# Nectar larceny in the trainbearers (*Lesbia*, Trochilidae)

**DOI:** 10.1101/2020.06.10.143909

**Authors:** Boris Igić, Ivory Nguyen, Phillip B. Fenberg

## Abstract

Many flower visitors engage in floral larceny, a suite of so-called ‘illegitimate’ visits in which foragers take nectar without providing pollination services. The data on prevalence of illegitimate visits among hummingbirds, as well as the total proportion of foraging and diet that such visits comprise is broadly lacking. Here, we report the occurrence of nectar larceny in both currently recognized species of trainbearers and analyze the proportion of plant visits categorized by mode of interaction as: primary robbing, secondary robbing, theft, and/or pollination. To the best of our knowledge, we provide the first published report identifying robbing in these species. We augment our original field observations using a trove of data from citizen science databases and literature. Although it is difficult to distinguish primary vs. secondary robbing and theft vs. pollination, we conservatively estimate that ca. 40% of the recorded nectar foraging visits involve nectar robbing. Males appear to engage in robbing marginally more than females, but further studies are necessary to confidently examine the multi-way interactions among sex, species, mode of visitation, and other factors. We discuss the significance of these findings in the context of recent developments in study of nectar foraging, larceny, and pollination from both avian and plant perspectives.

## INTRODUCTION

A growing list of bird species across several clades are known to forage for nectar on so-called ‘illegitimate’ flower visits, in which flower rewards are taken without the requisite provision of pollination services. Also termed ‘floral larceny,’ this mode of foraging is correspondingly gaining a broader appreciation as an important factor shaping the ecology and evolution of plant-animal interactions (Lara and Ornelas, 2001; Irwin et al., 2010; Rojas-Nossa et al., 2016; Boehm, 2018). Although species such as flower piercers (*Diglossa*, Passeriformes) are widely known to depend on nectar larceny, there are many reports of illegitimate visits by hummingbirds (Lara and Ornelas, 2001; Gonzalez and Loiselle, 2016). Some morphological characteristics of hummingbirds, including bill length and tomial serrations, are thought to be particularly closely associated with nectar larceny (Ornelas 1994, but see Rico-Guevara et al. 2019).

Nearly all plants that provide nectar and pollen rewards experience larceny, most often from insects and vertebrates, and many defend against it (Irwin et al., 2010). The remarkable frequency of illegitimate visits necessitated the development and adoption of a more precise lexicon of larceny (Inouye, 1980), which attempts to separate it into canonical modes, partly in service of conceptual clarification useful in pursuit of identifying its ecological and evolutionary causes and consequences. Thus, “primary nectar robbers” mechanically create a hole in flower tissue through which they remove nectar, bypassing the floral opening. By contrast, “secondary nectar robbers” remove nectar utilizing openings previously fashioned by primary robbers. Deceptively resembling pollinators, “nectar thieves” also access nectar through the floral opening. However, due to a mismatch between flower and thief morphology or behavior, pollination does not take place. Finally, some plant species are vulnerable to “base workers,” visitors that can probe along the base of the flower, between the free petals, and obtain nectar while bypassing both the anthers and/or stigma and the requisite damage or robbing.

These modes of nectar foraging may be difficult to distinguish from each other and/or legitimate visits— pollination—without careful observations and manipulations. Moreover, they can clearly quantitatively overlap so that, for example, thieving may merely reduce pollination efficiency without eliminating it, or individual birds can engage in a mix of primary and secondary robbing. While many studies report one or more modes of nectar foraging among hummingbirds (McDade and Kinsman, 1980; Roubik et al., 1985; Feinsinger et al., 1987; Ornelas, 1994), we are unaware of any published assessment of their relative importance, which is critical for a more complete understanding of plant-animal interactions, and the incentives and behaviors that drive the observed patterns.

Here, we present original observations and a larger meta-analysis of nectar foraging by Green- and Black-tailed Trainbearers (*Lesbia nuna* and *L. victoriae*), and discuss their significance in the context of plant-pollinator co-evolution and bill morphology. The original observations took place in and around Ollantaytambo, Peru, on *Brugmansia sanguinea*, *Fuchsia boliviensis*, and *Passiflora tripartita*. We coarsely document the proportion of pollination, theft, and primary- and secondary-robbing visits to flowers, by leveraging images from citizen-science databases eBird, iNaturalist, and other photographed/reported occurrences in the literature. Our highly preliminary analyses find that floral larceny is surprisingly common, at least ca. 40%, and that secondary robbing may be more common among hummingbirds than is generally appreciated, further complicating the proposed association between tomial serrations and mode of nectar foraging (Ornelas, 1994; Rico-Guevara et al., 2019).

## MATERIALS AND METHODS

### Species and site description

The genus *Lesbia* is currently comprised of two short-billed hummingbird species, Green-tailed Trainbearer, *Lesbia nuna* (Lesson, 1832), and Black-tailed Trainbearer, *Lesbia victoriae* (Bourcier & Mulsant, 1846). Their geographic distributions largely overlap, ranging from Colombia to Peru in montane scrub and other semi-open habitats. The species are differentiated by color, specifically the extent of emerald green on dorsal side and tail (more in *L. nuna*). Black-tailed Trainbearers are generally larger, as well. Both species are strongly sexually dimorphic, with males in possession of a spectacularly long tail. The species are difficult to differentiate and confidently assign because of a great amount of geographic variation within each species (many subspecies names exist) and possible existence of undescribed species (Weller and Schuchmann, 2004; Stiles, 2004). These difficulties are especially apparent in attempts to identify individuals from single photographs, and we consequently conducted our analyses (see below) at the generic level.

On January 14th, 2019, we recorded and photographed a male Green-tailed Trainbearer, *Lesbia nuna*, which nectar-robbed several flowers of *Fuchsia boliviensis* (Onagraceae) in Ollantaytambo, Peru (13°15′44″S, 72°16′14″W), and then followed the same individual on a 50m foraging bout, while it nectar-robbed flowers of *Brugmansia sanguinea* (Solanaceae)—a species with spectacular ca. 30cm-long tubular crimson-to-yellow flowers—and then legitimately visiting *Salvia leucantha* (Lamiaceae). Both rusty and black-throated flowerpiercer (*Diglossa sittoides* and *D. brunneiventris*) occur at relatively high densities in the area (B.I. and P.B.F., unpub. data), are known to pierce both *Fuchsia* and *Brugmansia* species, and may have been responsible for the previously made calyx and petal hole at the base of *B. sanguinea* flower.

### Literature searches

We examined the literature and internet resources to accumulate a database of additional instances of nectar collecting visits by *Lesbia* species. First, we used internet and Google Scholar exact keyword searches of the existing scientific literature, including: ‘Datura’ or ‘Brugmansia’ and ‘Lesbia’ or ‘trainbearer’; ‘Lesbia’ and ‘diet’. We employed “advanced search” filtering to reduce the considerable volume of inappropriate content. Documentation of any details on foraging ecology of the trainbearers is sparse. We found two references hinting at illegitimate flower visits by trainbearers. Gould (1861, p.15) cited in Ornelas (1994), contains a passing reference to piercing of a *Brugmansia sp.* flower by a *Lesbia sp.*, but the visit was inferred as insect predation, not nectar robbing, as later re-framed by Ornelas (1994), and neither the species nor locations were identified. In a paper documenting secondary robbing behavior in Cinereous Conebill, Vogt (2006) mentions unpublished observations of secondary nectar robbing by a trainbearer. Our literature search and two substantial meta-analyses (Ornelas, 1994; Irwin et al., 2010) did not retrieve any other references to illegitimate visits by trainbearers. Additionally, although we could find no papers documenting their foraging ecology in any detail, gut contents of a single individual examined did include arthropods (Remsen et al., 1986). Therefore, as generally holds for hummingbirds, the trainbearers’ diet minimally contains arthropods and nectar. The mode of nectar foraging is, however, unclear.

### Foraging data collection and scoring procedures

Consequently, we examined all identified species records for the Genus *Lesbia* in two public databases— iNaturalist (iNaturalist, 2019) and eBird (Sullivan et al., 2009), and on the online photo service flickr (https://www.flickr.com). For all resulting records, we applied the same methodology. Images without any flowers were discarded, as were those in which birds were simply perched or flying near a flower. We then closely examined the remaining images, in which birds were hovering in close proximity (about a body length) to flowers, and facing them. For that subset, we extracted the location, date, and comment metadata. We scored individual sex (‘m’,‘f’, or ‘NA’), species identification (scoring specific epithets as ‘nuna’, ‘victoriae’, or ‘sp.’), estimated mode of interaction (primary nectar-robbing, secondary nectar-robbing, thieving, pollination, or ‘NA’), as well as our confidence in the estimate of the mode of interaction (‘low’ vs. ‘high’; to help facilitate any re-examination of our results). Finally, with the help of several colleagues, we identified the visited plant species.

Often, modes of interaction are unclear and we combined terms for the most accurate characterization of interaction. For example, primary vs. secondary nectar robbing and pollination vs. thieving are generally hard to distinguish, so we simply combined the likely interactions. This enables us to conservatively summarize modes of interactions as approximate ranges, without making unnecessary (and incorrect) assumptions. We should note here that nearly all of the combined pollination/thieving visits appear to have been legitimate visits, but establishing the extent of pollen transfer is prohibitively demanding for most studies of modes of nectar feeding. Similarly, we recorded the species identifications, but found them to be implausible or uncertain for a number of individuals, and we consequently highlight the jointly presented feeding habits for all of *Lesbia*. Because of these limitations, as well as the fact that the observations were not randomized in any way, we simply report a number of pooled summary statistics. Nevertheless, pooling over the ignored variable(s) can be misleading, so we perform limited three-way visual analyses, but caution against their over-interpretation. We coarsely examine associations between species ID (nuna/victoriae), sex (f/m), and binarized mode of interaction with flowers (robbing/other), examining the null hypothesis that the three traits are mutually independent. For model selection, we used log-linear models implemented in R package MASS (R Core Team, 2019), and AIC scores from LRstats. The nature of our data collection and resulting exploratory analyses suggests that any associations ought to be re-tested by obtaining a new set of data in advance of further testing.

### Bill morphology

Short-billed hummingbirds and those with serrated tomia are thought to be more likely to nectar-rob flowers with long corolla tubes, co-adapted with long-billed hummingbirds (Lara and Ornelas, 2001). We did not find a reference to presence/absence of serrated tomia in trainbearers, so we scored this trait in 17 specimens of *Lesbia nuna* and four specimens of *Lesbia victoriae* at the Field Museum of Natural History. We used a dissecting microscope to visualize and score this trait on the available collections, as well as measure their bill lengths (tip to operculum).

All accessions, metadata, image locations, and scored data used in our analyses are available online in Supplemental Materials.

## RESULTS

### Field observation of robbing

In our field observations, we recorded a single adult male foraging bout, during which this individual visited flowers of three plant species. Approximately ten flowers of *Fuchsia boliviensis* were robbed (unclear if primary or secondary robbing took place) as well as two flowers of *Brugmansia sanguinea*, while five flowers of *Salvia boliviensis* were apparently legitimately visited (ruling out or confirming pollination success requires controlled experiments). Photographs collected in the course of observations were deposited in iNaturalist (https://www.inaturalist.org/observations/37206000; iNaturalist 2019).

### Database collection

We collected and examined more than 1400 photographs of *Lesbia* and found floral visits for 180 foraging bouts (Figure 1, and Supplementary Data Table) on at least 42 flowering plant genera, identified thus far. It is unlikely that many of these instances are replicates, recording the same individual, although this is somewhat uncertain for the vicinity of Cerro de Monserrate in Bogota, Colombia, with about a dozen documented foraging bouts.

We assigned the mode of interaction for 135 visits with some confidence, while the mode remains unknown for 45 visits, most often because pictured birds were hovering in the vicinity of a flower, immediately before or after nectaring. The most common recorded mode was ‘pollination/thieving’ (46 visits), a category that we scored conservatively (vaguely) because it is clearly continuous, virtually impossible to confidently score separately without functional study. Ideally, one would examine pollen transfer or effected seed set, after bagging and marking flowers. Our impression, however, is that the majority of these visits at least plausibly resulted in pollen transfer. The second most common mode was combined ‘nectar robbing’ (primary and secondary robbing; 41 visits) indicating that an individual fed through a pierced hole in the side of the corolla. It was followed by ‘pollination’ (36 visits), which indicated a fairly clear legitimate interaction, ‘secondary nectar robbing’ (10 visits), and thieving (2 visits). Although we could not confidently assign any visits to the ‘primary robbing’ mode, this is entirely due to the one key limitation of our observational approach, reliance on scoring from photographs. If we conservatively lump all ‘pollination/thieving’ and ‘pollination’ observations as legitimate, such visits comprise approximately 60.7% of the total and illegitimate visits 39.3% (combining all robbing and thieving).

**Figure 1.**
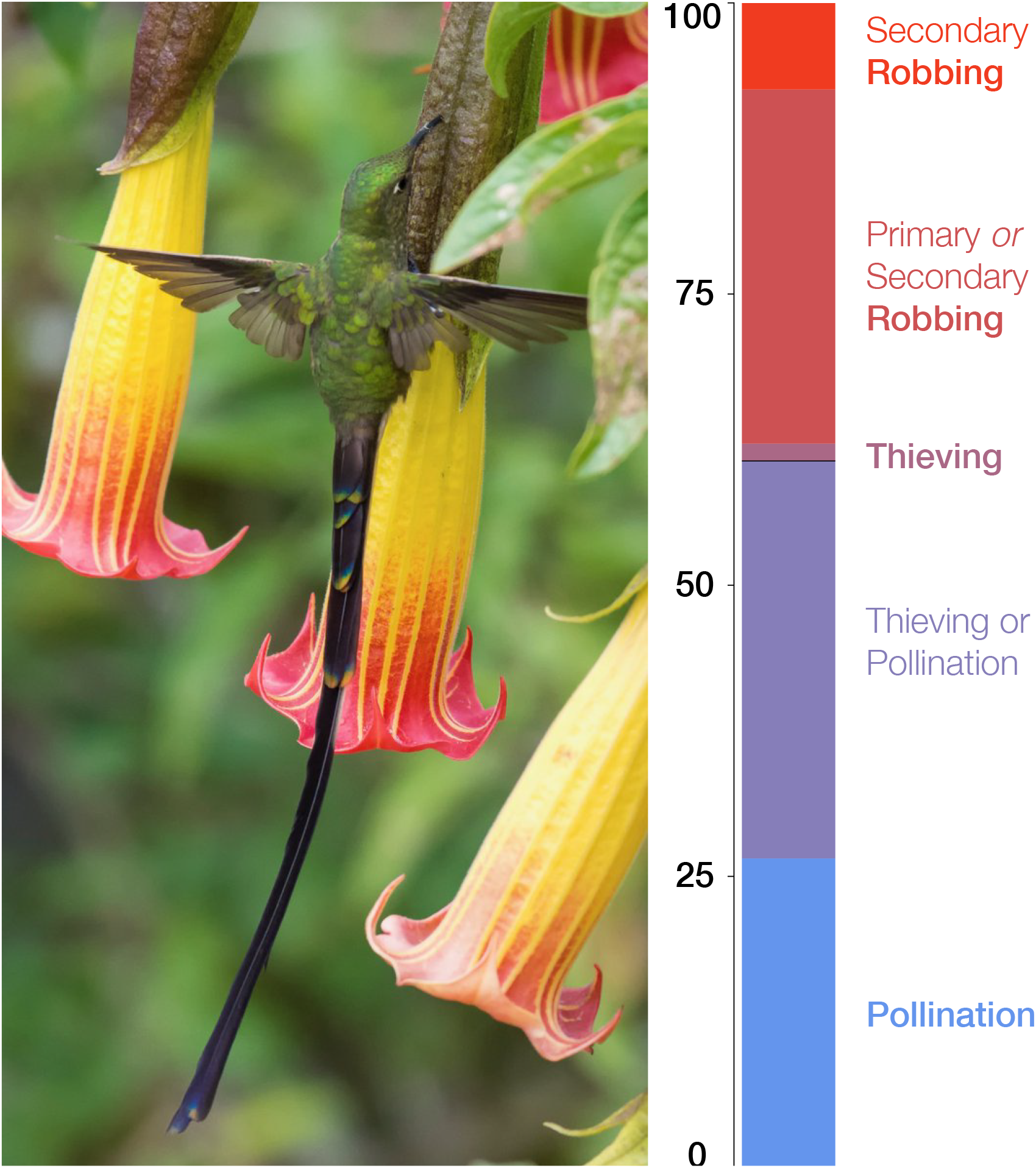
A nectar robbing visit by a Black-tailed Trainbearer *Lesbia victoriae* on *Brugmansia sanguinea*, whose flower was previously pierced by a species of *Diglossa* (Photo by Diego Emerson Torres; used with permission). Right panel: A bar plot illustrating the relative visit mode frequencies of *Lesbia* species (n=135). Of the 135 visits whose mode is known (could be assigned), 60.7% were legitimate (combining pollination and pollination/thieving) and 39.3% illegitimate (combining all robbing and thieving).

With a number of limitations in mind, we present a few preliminary analyses in an attempt to estimate the association of species ID, sex, and mode of visitation. When these factors are jointly considered in a three-way contingency table, we find that three variables are likely not mutually independent (AIC=56.43, LR *χ*^2^=13.55, d.f.=4, p¡0.01). Non-robbing visits by *Lesbia nuna* females and robbing visits by *Lesbia victoriae* males have the highest residuals. The lowest AIC model posits conditional independence (mode is independent of species, given sex; AIC=48.78, LR *χ*^2^=1.91, d.f.=2, p=0.39), indicating that sex differences are greater than species differences. One broad interpretation is that the females generally rob less than the (territorial) males. We emphasize that this result is preliminary, and the associations may stem from a number of data limitations, especially confounding unmeasured variables, or those not considered *a priori*. For example, more *Lesbia victoriae* males were observed in urban parks where plant communities and robbing frequency may differ, compared to rural or natural areas. Moreover, a model with all two-way interactions, but without the three-way interaction of the saturated model, provides only slightly worse fit (ΔAIC=0.15). It is therefore presently unclear whether whether species or sex differences are more important in predicting nectar foraging modes.

### Bill morphology

We examined 17 specimens of *Lesbia nuna* and found no serrated tomia (Figure 2). The imbalance of sex ratio (15 males) is concerning, but perhaps unimportant in this case, because serrations are more likely to be present in males (Rico-Guevara et al., 2019). The mean bill length for males was 15.9 ± 0.7 mm, and for females 14.2 ± 3.2 mm. Similarly, we found no tomial serrations in four specimens (three male, one female) of *Lesbia victoriae*. Their mean bill length for males was 14.7 ± 0.4 mm, and one female measured 11.0 mm.

**Figure 2.**
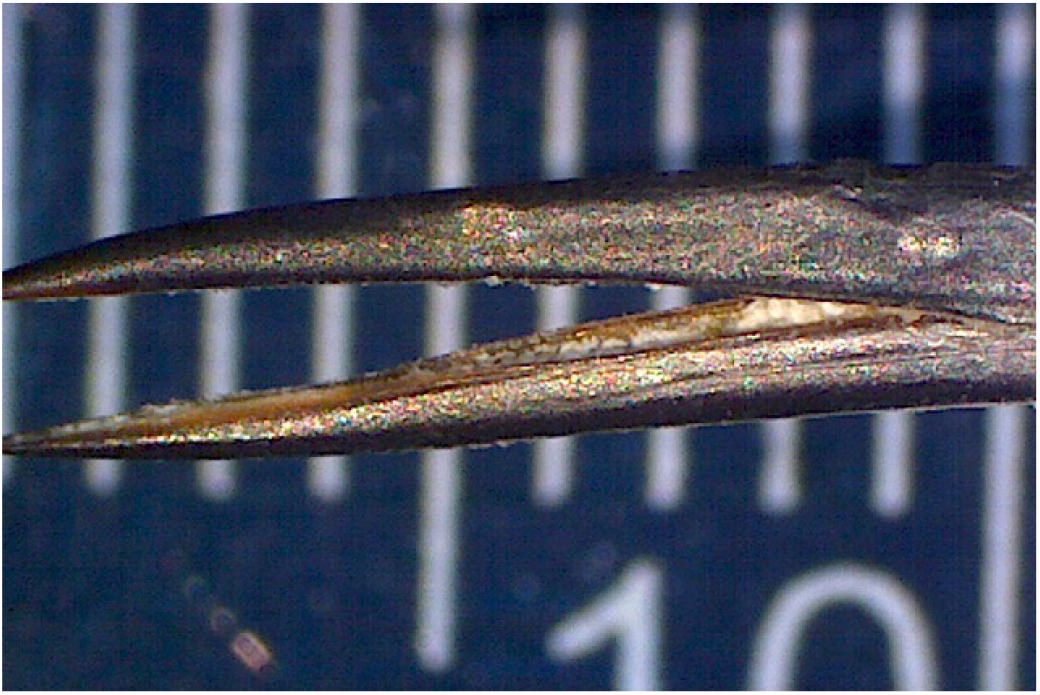
Anterior side of the bill of a male *Lesbia nuna*, FMNH-222250, with no conspicuous serrations on its mandibular (or maxillary) tomia. For scale, a ruler denoted in millimeters is seen in the background.

## DISCUSSION

Hummingbirds are generally thought to serve as legitimate flower visitors, and effect pollination in exchange for host plant nectar. There is, however, an increasing appreciation of the fact that nectar obtained during illegitimate visits—including both primary and secondary robbing, as well as thieving— may comprise a substantial portion of diets of many species (Lara and Ornelas, 2001; Irwin et al., 2010; Boehm, 2018). Presently, the frequency distribution of nectar larceny within and among bird individuals and species, and a variety of covariates (community composition, territoriality, seasonality, geography, etc.) is unclear. Most reports of hummingbird nectar foraging document the presence (or imply absence) of nectar robbing, and do not provide an estimate of the proportion of visits dedicated to illegitimate visits. This is in large part due to the difficulty of accruing this kind of data and, when larceny occurs, separating its modes.

Our study originated with a simple natural history observation of robbing by a trainbearer, which we augmented using a trove of citizen-science data from crowd-sourced public databases. We believe that such estimates are a first step in improving our understanding of the dynamics between plants and pollinators, especially why and how (e.g. primary vs. secondary robbing) some individual birds or species steal more than others, how larceny affects plants, and how they respond to larceny. Below, we discuss the results and their immediate implications of this small-scale study and place them in the context of the literature on ecology and evolution of plant-hummingbird interactions.

We find that both currently described species of *Lesbia* steal nectar, mostly by robbing. Unambiguous illegitimate visitation constitutes a substantial proportion, around 40% of all nectar foraging visits. A few previous studies indicate that, for the species surveyed at the individual-plant level, the frequency of nectar robbing by birds on plant species can be very high (although variable; Arizmendi 2001). An unpublished report (D. Boose, cited in Maloof and Inouye 2000) found that robbing may comprise up to 92% of all Stripe-tailed Hummingbird (*Eupherusa exima*) visits in *Razisea spicata*. Primary and secondary robbing are generally difficult to disentangle from casual observations (Vogt, 2006; Boehm, 2018). Confidently scoring the specific mode of robbing (primary vs. secondary) demands evidence bearing on the question whether ‘breaking-and-entering’ took place, which in turn requires careful experimental data collection that we could not directly collect. For example, capturing the act of piercing of flowers or documenting entry through a pre-existing hole could enable unambiguous scoring of robbing mode. We find that secondary nectar robbing is relatively common, comprising between 7.4% and 37.8% of the visits (Figure 1). This large range estimate reflects uncertainty (not variability) due to the limitations of our approach. However, we examined additional photographs of many robbed plant species, which frequently show extensive flower damage, and found records of several species of flowerpiercers (*Diglossa*) commonly observed in the areas in close proximity to trainbearer robbing locations (B.I. and P.F. unpub. data). It is therefore likely that secondary robbing may represent the majority of illegitimate visits by trainbearers. Moreover, it is possible that this mode of nectar foraging is underestimated in other hummingbird species.

From the hummingbird perspective, if primary robbing is more likely to allow access to previously unvisited, relatively nectar-rich flowers, then perhaps secondary robbing ought to be less advantageous (Irwin et al., 2010). Framed in this way, any secondary robbing may appear somewhat paradoxical. It is possible that, instead, one or more of the common underlying assumptions are flawed. Specifically, we can imagine that, for example, if they are faced with resource limitation (of flowers that are ordinarily legitimately exploited), trainbearers respond by robbing or thieving from inferior resources, a situation that may be exacerbated by territoriality. Although our study cannot strongly substantiate it, a trend of higher robbing frequency in males than females (complicated by association with species), may be related to such phenomena. On the other hand, the evolutionary game between hummingbirds, flowers, and all other interactors (whole pollinator guilds, herbivores, etc.) is likely substantially more complex than is generally appreciated (e.g. Maloof and Inouye 2000), broadening the set of conditions for stable non-zero frequencies of secondary robbing.

Floral larceny is often targeted at flowers of plants that have close co-evolutionary relationships with other legitimately pollinated animal species, such as the sword-billed hummingbird and its guild of long-flowered species (Soteras et al., 2018). Some characteristics of hummingbirds, including bill length and tomial serrations are thought to be particularly closely associated with primary nectar robbing (Ornelas, 1994). The hypothesis that the latter trait, serration of tomia, affects nectar robbing ability and efficiency remains broadly untested. Recent studies suggest that the presence of serrations is sexually dimorphic, with greater or exclusive expression in males, and that this feature could instead be shaped by a role in plumage preening or fighting, especially male territorial defense (Rico-Guevara et al., 2019). Although additional field studies of nectar robbing and examination of bill morphologies for the presence of serrate tomia are needed to clarify their role, our data provide some insight. First, if secondary nectar robbing is more common than previously thought, the association between serration and (all) robbing observations may be lower and more difficult to detect. Serrations may well provide flower-robbing utility, but they are clearly unnecessary for secondary robbing. Second, if males rob more frequently, then the existence of sexual dimorphism for tomial serrations is not problematic for the Ornelas hypothesis. Moreover, neither sexual dimorphism nor the proposed role in defense precludes simultaneous or exaptive function of serrated tomia in piercing. Consequently, it will be difficult to accumulate a great weight of evidence that exclusively support or detracts from this hypothesis. Nevertheless, it is instructive that many other specialized piercers, such as the Wedge-billed Hummingbird (*Schistes geoffroyi* and Purple-crowned fairy *Heliothryx barroti*, fail to present sexually dimorphic serrations (Rico-Guevara et al., 2019). A systematic search of nectar feeding mode, along with a number of predictors, and coupled with careful studies of feeding behavior and morphology, are likely to be most persuasive in examining the validity and generality of the Ornelas hypothesis.

Predominance of different modes of larceny is likely to be associated with differing evolutionary responses from flowering plants, including variation in amount and/or time-dependence of nectar release, mating system (increased reliance on self-fertilization), and defenses, both physical (spines, thicker calyces or bracts) and chemical (secondary metabolites). A number of plant species visited by trainbearers are members of the nightshade family (Solanaceae), known for common production of alkaloids, which are broadly toxic to vertebrates. Perhaps not coincidentally, a great number and quantity of alkaloids are produced by species of *Brugmansia* and *Nicotiana*, and tissue at the base of flowers (calyx) contains some of the highest concentrations of such compounds (Saitoh et al., 1985; Alves et al., 2007). *Brugmansia* species have not been studied in equal detail to tobacco, but at least some plants in this genus can incapacitate juvenile humans, orders of magnitude larger than hummingbirds, following casual skin contact (Andreola et al., 2008). Moreover, given that there is a strong association between hummingbird pollination and self-fertility (Wolowski et al., 2013), stereotypically hummingbird-pollinated flowers may be more commonly robbed by both avian and hymenopteran robbers. This may be especially true for those flowers with relatively longer tubular corollas and large amounts of nectar (Lara and Ornelas, 2001; Rojas-Nossa et al., 2016), precisely the syndrome commonly associated with hummingbird pollination.

Our observations and analyses point to additional needed work on systematics and ecology of *Lesbia*, especially their unsettled taxonomy, feeding ecology and energy budgets. More broadly, we sorely lack observational data to establish the relative frequencies of nectar feeding modes and manipulative field study to obtain direct evidence for nectar feeding dynamics across all hummingbirds and other nectar-feeding birds (Irwin et al., 2010). Our study documents the presence of larceny in trainbearers and, more specifically, shows that floral larceny is surprisingly common. These findings indicate that larceny, especially secondary robbing, may be more common among hummingbirds than is generally appreciated. Furthermore, we demonstrate that citizen science databases can be fruitfully employed to establish the presence of nectar robbing and coarsely examine its frequency. Such approaches can also generate hypotheses and guide an improved understanding of plant-side effects and responses, including pollination efficiency, the distribution of visitor interactions (Maloof and Inouye, 2000; Arizmendi, 2001), plant incentives and defenses (as well as overall fitness) (Pelayo et al. 2011), and ultimately better explain the dynamics in the co-evolutionary game between flowers and their visitors.

**Table 1.**
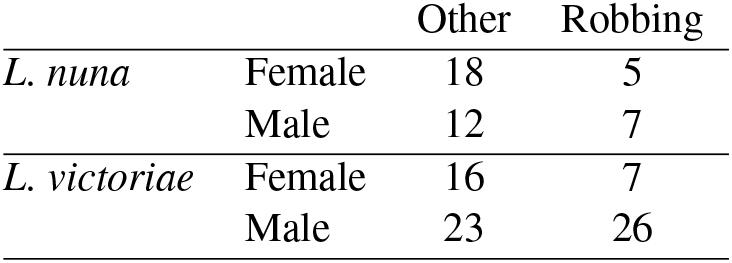
Cross tabulation data of nectar foraging mode, sex, and species identity for *Lesbia spp.*. ‘Other’ denotes lumped pollination and thieving, and ‘Larceny’ combines both modes of robbery. Sex identification relied on tail length and subtle color differences as a proxy, and species identification was in nearly all cases copied from the original source. Of 180 total foraging bouts captured, we assigned visit mode to 135, and each of visit (nectar foraging) mode, sex, and species to 114 shown in the table.

## ACKNOWLEDGEMENTS

We thank iNaturalist and eBird projects, and all contributing citizen naturalists whose images and data enabled us to conduct this study. We thank Diego Emerson Torres (https://ebird.org/profile/NzU3NjIy) for permission to use the image in Figure 1. C. Rushworth and C. Rothfels provided generous help with plant species identification, and J. Treehorn for fixing the cable. This work was supported in part by the National Science Foundation NSF DEB-1655692.

---

**Supplementary Figure S1.**
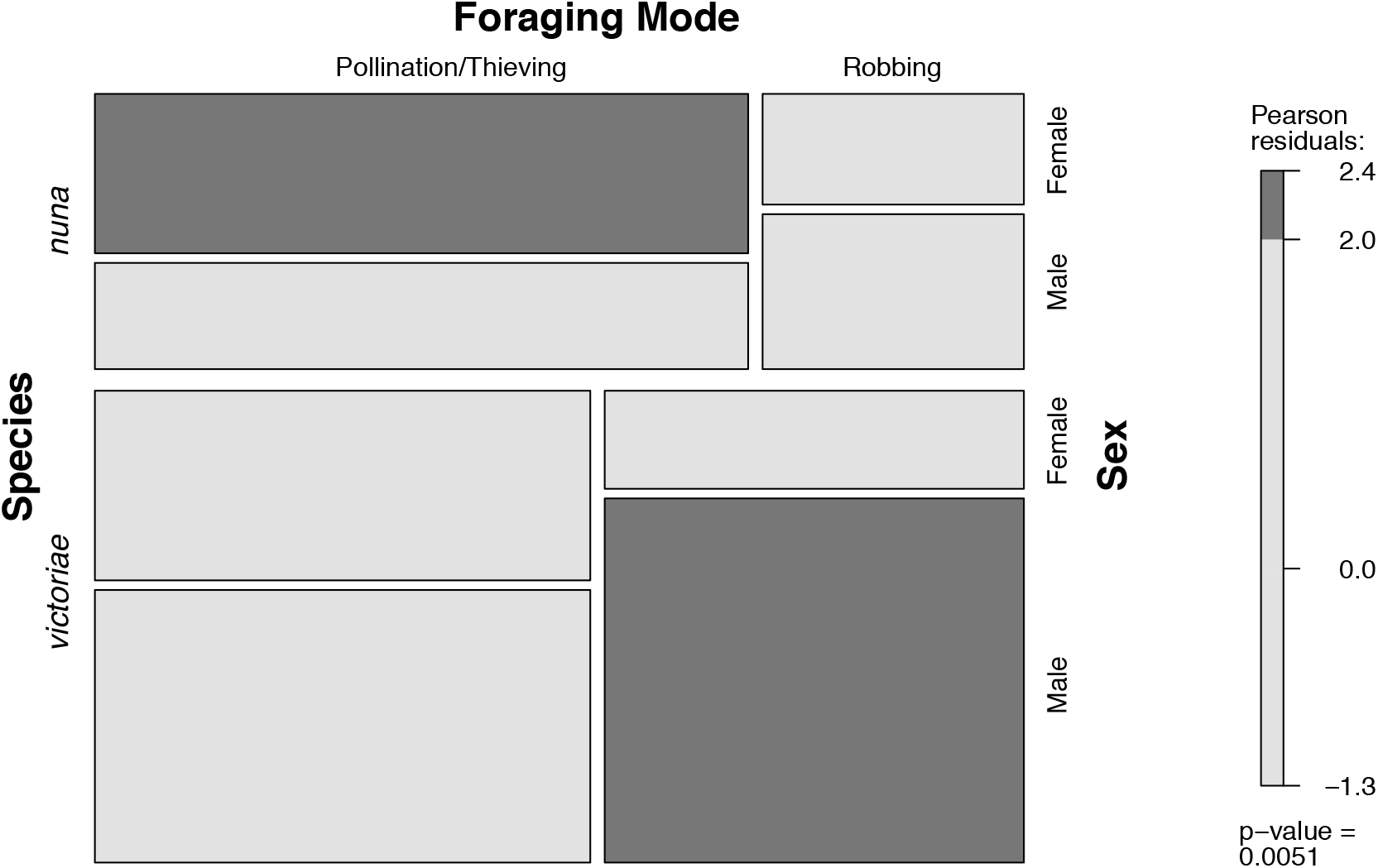
Mosaic plot of data presented in Table 1, illustrating the three-way contingency table. Cells sizes are proportional to observation number for each category. Shading color illustrates values of Pearson residuals (dark = greater than 2.0), as shown in scale on right.

## Notes

### Competing Interest Statement

The authors have declared no competing interest.

